# Interactions between discrete events and continuous dynamics in the regulation of scallops valve opening: insights from a biophysical model

**DOI:** 10.1101/2020.12.25.424408

**Authors:** Jean-Marc Guarini, Jennifer Coston-Guarini, Luc A. Comeau

## Abstract

This study constitutes a first attempt to quantify processes that govern valve gape dynamics in bivalves. We elected to focus on the scallop, *Pecten maximus*, not only because of its economic importance but also because it has a complex behaviour and high sensitivity to stress, which can be inferred from valve gape dynamics. The adductor muscle is the primary organ implicated in valve movements. Scallops, as other bivalves, move their valves sharply to ensure basic physiological functions or to respond to stressing conditions; these sharp events can be perceived as discrete events within a continuous dynamic. A biophysical model, originally designed for human muscles, was first selected to simulate the adductor muscle contraction, countering the passive valve opening by the umbo ligament. However, to maintain the possibility of rapid valve movements, described as typical of bivalves behaviour, it was necessary to modify the model and propose an original formulation. The resulting hybrid modelling simulates how valve opening tends to converge continuously toward a stable steady-state angle, while being interspersed with discrete, sharp closing events, deviating values from this equilibrium. The parameters of the new model were estimated by optimization using Hall-Effect Sensor valvometry data recorded in controlled conditions. Equilibrium of the continuous regime (when fiber activation equals deactivation) was estimated for a gape angle close to *ca*. 15 degrees, which is *ca*. 45% of the maximum opening angle, hence implying a constant effort produced by the adductor muscle. The distribution of time intervals between two successive discrete events did not differ significantly from a random process, but the peak amplitudes deviated from randomness, suggesting they are regulated physiologically. These results suggest that discrete events interact with continuous dynamic regimes, regulating valve opening to minimize physiological efforts and conserve energy. However, because the overall physiological state of the scallop organism conditions the activity of the adductor muscle, a complete understanding of the physiology of bivalves will require linking a more comprehensive model of valve gape dynamics with experimental observations of physiological energy consumption under different conditions.

## 1 Introduction

Bivalves open their shells to perform essential physiological functions as respiration, feeding, reproduction, and feces expulsion. Bivalve ”valvometry” (Marceau 1909; Rao 1954) was developed over a century as a means to monitor this activity and infer behaviors under different environmental conditions (see reviews by Kramer et al. 1989; Clements and Comeau 2019). Valvometry became a generic term which encompasses many different techniques; it started with “sooted glass” techniques (Marceau 1909), and gained automation with strain gauges (Wilkens 1981), Hall-effect sensors (HES) (Nagai et al. 2006), impedance electrodes (Tran et al. 2003), and fiber optic sensors (Franck et al. 2007). Because bivalves close their shell in response to stress, valvometry has been applied not only to study their physiology (Payton et al. 2017; Comeau et al. 2018) but also to characterize responses to environmental perturbations (Gaine and Shumway 1988; Nagai et al. 2006, Redmond et al. 2017). Valvometers based on HES have many advantages over other techinques: HES are light, compact and require only one connection to collect data. However, they require careful implementation to ensure a suitable estimate precision; an *ad hoc* calibration based on both electromagnetic properties of the sensors and dynamic geometry of the shell was developed (Guarini et al. 2020) to obtain time series and dynamics indicators of valve gape variations; it permitted comparisons between data series from individual organisms.

Closure of the two valves (shells) occurs in response to contractions of the adductor muscle(s). This contraction opposes a continuous opening force generated by a ligament located at the umbo (Bayliss et al. 1930; Trueman 1953). The adductor muscle itself has two parts characterized by two different types of fibers. The large, striated part of the muscle is responsible for fast movements, while the smaller, smooth muscle fiber is responsible for slower, sustained and cyclic movements. Wilkens (1981) suggested that this sustained smooth muscle tension exerted against the force of the hinge (umbo) ligament is the explanation for the half-open position of the valves described as a resting posture. To the best of our knowledge, however, there are no quantitative studies linking valve gape dynamics to muscle activity, even if there are many studies on the adductor muscle in bivalves, and especially in scallop species (see Chandler 2006 for a review).

In fact, the vast majority of modelling studies about muscles concern humans. In the late 1930s, Hill (1938) proposed a first quantification of human muscle dynamics from a bioenergetics point of view. Currently there are three types of models developed for human physiological studies: biophysical, mechanical and biochemical (Ruina 2016). All three are based on a common principle that muscle fibers are excited by the nervous system and this induces the release of Ca^2+^ cations that modify the protein configurations producing contractions and generating force. These models address the problem of muscle fatigue which results from a decrease in muscle fiber activation and contraction, and a decrease in the force developed by the muscle. In these models, the relaxation of the muscle after contraction can be considered either a passive process or active deactivation process in these models.

Our global objective is to establish a quantitative framework for interpreting physiological signals obtained from king scallops (*Pecten maximus*) with valvometry. In the present study, a minimal biophysical system was designed to represent the adductor muscle dynamics and simulates activation, deactivation, and recuperation processes. This submodel was then used to calculate the forces applied by the muscle on each valve.

To minimalize the system of equations, the model was conceived to describe a global adductor muscle functioning, without distinction between smooth and striated parts. In addition, only continuous cahnges interspersed with single events were investigated (as opposed to series of reflex contractions). The model was used to test hypotheses regarding the underlying processes that govern the passive action of the ligament and the active reactions of the adductor muscle. For achieving these objective, in this article, model outputs were compared with data collected on scallop individuals equipped with HES valvometers in tank experiments. Once HES were calibrated with the method we developed in an earlier article (Guarini et al. 2020), the comparison between time series and simulations were achieved by optimization, allowing unknown parameters to be identified. We then describe statistically robust basis for interpreting *in situ* valvometry time series, including those for which individuals are affected by environmental stressors (e.g., temperature change, eutrophication, harmful algal blooms). General pespectives were drawn to develop further the model for two different research directions, ecophysiology of bivalves, their monitoring in aquaculture context, and for extending their use as sentinel species in environmental impact assessment applications.

## 2 Material and Methods

### 2.1 Modeling the dynamics of the angle of the shell, *α* (*t*)

To formulate our model, we introduced an angle, *α* (in radian) formed by two rays placed on each valve in such a way that *α* = 0 when the shell is closed (see Figure 1). The origin (i.e. the angle vertex) is located at the shell umbo, and the plan formed by the two rays is perpendicular to the axis formed by the hinge. The dynamic of *α*(*t*) was calculated by integrating the angular velocity, v(t), which results from two forces; the first one is a passive force, *F_O_* (in Newtons), produced by the tension of a ligament, which opens the shell. The second one is an active force, *F_C_* (in Newtons), produced by the adductor muscle, which closes the shell.

**Figure 1:**
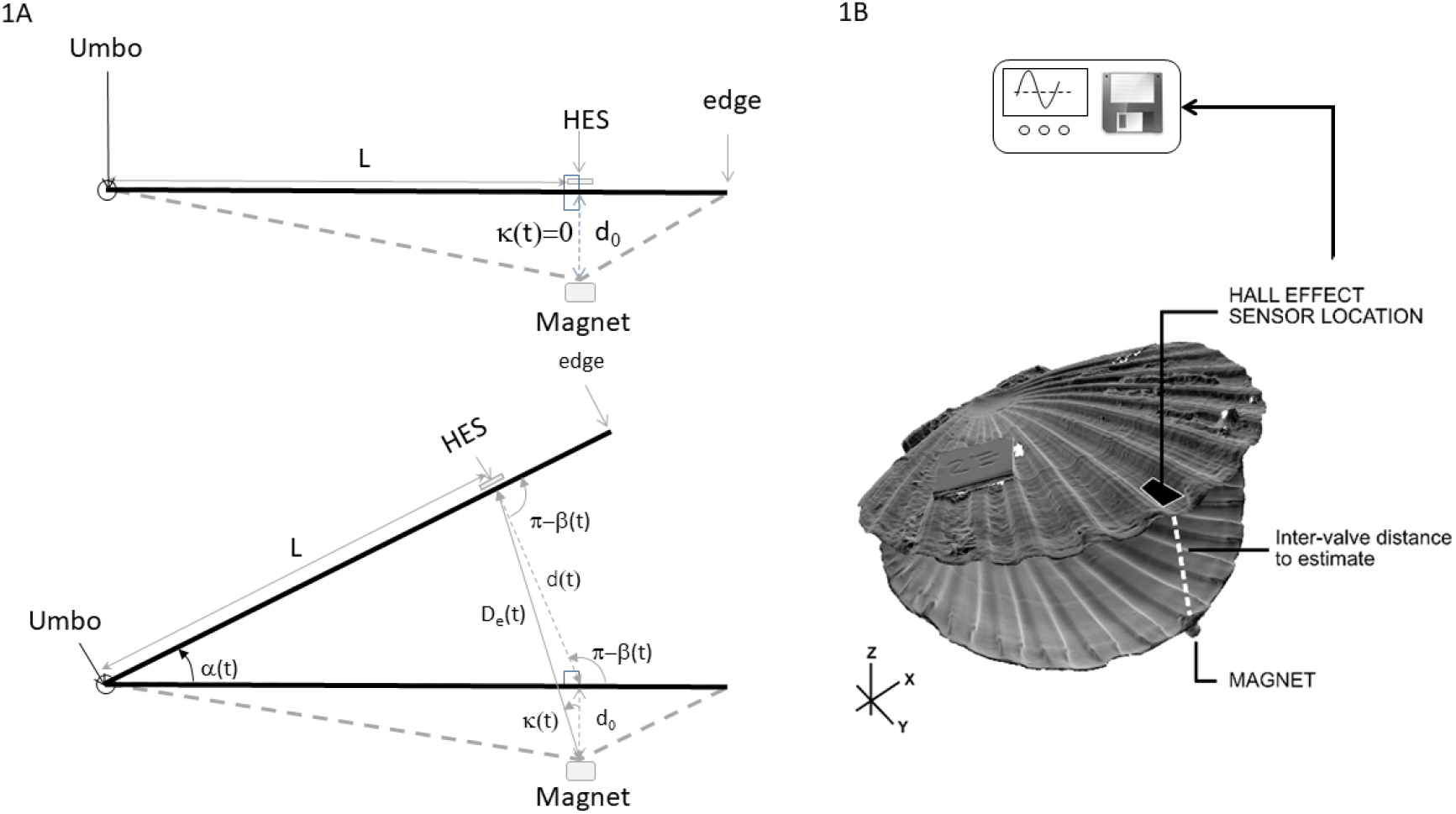
Hall-Effect experimental setup (A) Geometric representation of a cross-section of a *Pecten maximus* shell showing the point of placement of the Hall-effect sensor (HES) on the flat upper valve and the magnet on the curved lower valve. (B) Schematic of a scallop (*Pecten maximus*) shell showing the placement of the Hall-effect sensor (HES) on the flat valve and the magnet on the curved valve.

The module of the passive opening force, *F_O_*, which is an elastic restoring force, was considered to be a linear function of the ligament elongation:

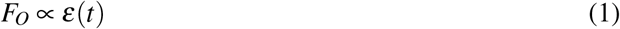

where *ε* (*t*) (dimensionless) represents the relative elongation of the ligament, function of the relative distance of the opening, and expressed as:

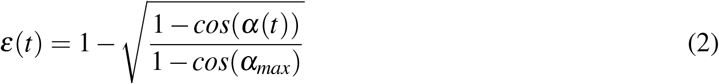

where *α_max_* (in radians) is the maximum opening angle which can be determined experimentally from the maximum opening distance *d_max_* (in mm). The relative elongation ranges from 0 (when *α*(*t*) = *α_max_* and the shell is fully open) to 1 (when *α*(*t*) = 0 and the shell is completely closed). The positions of ‘fully open’ or ‘fully closed’ are functional definitions from the calibration procedure (Guarini et al. 2020, see also section 2.3). Angular speed is determined by restoring forces applied and hence is linked to the quantity of kinetic energy produced. Therefore, the angular speed was expressed as a function of square of the elongation:

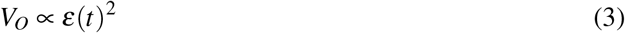

Conversely, the adductor muscle contraction induces a force, which counters the passive opening force by mobilizing muscle fibers. We assumed that this closing force, *F_C_*, is a function of the proportion between the number of activated muscle fibers, *m_A_*, and the total number of muscle fibers, *m_T_*:

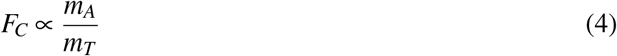

The total number of muscle fibers, *m_T_* is the sum of three components: *m_A_*, *m_F_* (the number of deactivated fibers), and *m_U_* (the number of inactive fibers which can still be mobilized). *m_T_*, hence the 3 components, depends on the size of the muscle. To get rid of this dependency, *m_A_*, *m_F_*, and *m_U_* were divided by *m_T_* to express the state of the muscle in terms of dimensionless proportions *p_A_*, *p_F_*, and *p_U_*, respectively. It has been proposed that the dynamics of the muscle can be simulated by a biophysical model (Liu et al. 2002; Böl et al. 2011; Looft et al. 2018):

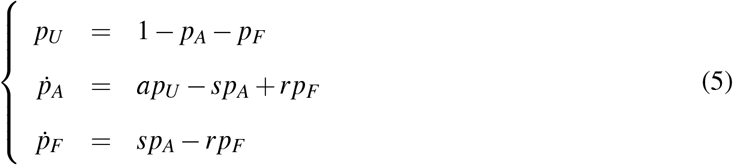

where a is the activation rate, s is the deactivation rate, and r is the recovery rate (all in *t*^−1^). According to Hill (1938), the speed of muscle shortening due to contraction of muscle fibers is a linear function of the closing speed:

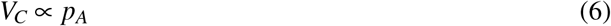

Thus, the resulting dynamics of angle opening, *α* (*t*), combining equations and [3] and [6], was expressed as:

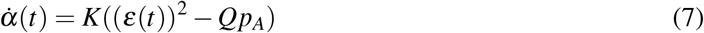

where K and Q are two parameters which need to be estimated. It remains difficult to provide a physiological, or even a mechanistic, meaning for these two parameters. However, from a mathematical point of view, K, has a dimension of angular velocity (*radian.t*^−1^), and Q (dimensionless) represents a scaling factor between two dimensionless variables: 1) the relative ligament elongation and 2) the proportion of active fibers in the adductor muscle.

### 2.2 Simulation of valve-gape angle dynamics

Shell valve movements are characterized by alternating series of opening and closing events. The opening event is considered to be passive and it does not require muscle fibers to be activated. It can be assumed that the organism controls the opening event by decreasing the activation rate, a. In contrast, closing of the shell is an active event, and activation of available muscle fibers is required. For fast closing events (identified by discrete closing peaks), we hypothesized that all available fibers (represented by *p_U_*) are mobilized at once to close the shell. The corresponding discrete transition system of this model was represented by:

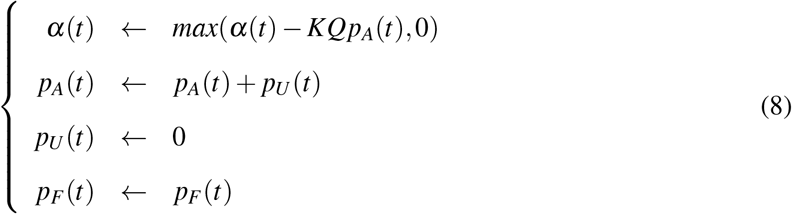

### 2.3 Data acquisition and calibration of sensors

Experiments were conducted on king scallops (*Pecten maximus*) in April 2018. Twelve *Pecten maximus* individuals were collected in the Bay of Brest (Western Brittany, France) at depths between 10 and 15 m. Their size were close to each other (their average maximum length, from umbo to the edge, was equal to 10.18 ± 0.25 (SE) cm). Shells were cleaned of epibionts, and were placed in holding tanks (7.3 x 6.2 x 4.2 *dm*^3^). Water in the tanks was continuously renewed at a constant flow rate of 15 *dm*^3^.*h*^−1^ with filtered seawater from the Bay. During this period, the scallops received a daily suspension of *Isochrisis galbana* (ca. 1L.*h*^−1^ at ca. 17.10^9^ cells.*L*^−1^). Artificial light was produced by light emitting diodes at a color temperature of 4500 K and with a periodic cycle of 12h light (from 8:00 am to 8:00 pm) and 12h dark. After a two-week acclimation period, scallop individuals were equipped with a Hall Effect Sensor (HW-300a, Asahi Kasei, Japan) according to a protocol designed by Wilson et al. (2005). One HES, sealed in epoxy resin, was glued along the axis of maximum length, in a position close to the edge of the flat valve; the average distance between the HES and umbo was 9.55 ± 0.25 (SE) cm. The HES was then connected to a data logger (Smart Dynamic Strain Recorder, DC204R, Tokyo Sokki Kenkyujo Company, Japan). A small neodymium magnet (diameter, 4.8 mm; thickness, 0.8 mm) was glued on the opposite curved valve surface directly below the sensor. Hall voltage was checked while the shell was still closed; a second magnet was added when needed, to set the Hall voltage closer to the maximum value. Total handling time was less than 25 min. Two sets of six scallops were placed in experimental tanks with filtered circulating sea water and the Hall voltages recorded at a frequency of 10 Hz for 7 – 8 days. At the end of the experiment, a calibration curve was generated for each scallop by measuring the voltage produced at 14 different known inter-valve distance values, after the adductor muscle was severed. This is done by inserting a series of glass wedges between the two valves at the point farthest from the umbo. Finally, HES measurements were recorded when the shells were fully closed and fully open. These data were used to create a calibration curve for each individual scallop. Both the experiments and the data collected are also described in an earlier article (Guarini et al. 2020)

The measurement of the Hall-effect voltage is proportional to the magnetic field intensity, B, which varies as a function of the inverse of the square of the distance between the HES and the magnet, *D_e_* (in mm) and, because of the rotation of the shell around the hinge, is a function of *κ* the angle between the axis of the magnetic field and the axis between the HES and the magnet surface (Figure 1). Therefore, the distance between the HES and the magnet was calculated as:

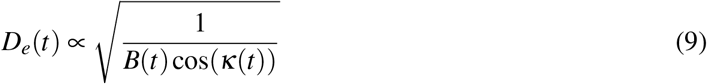

The intervalve distance, *d*(*t*) (in mm), was calculated from the estimates of *D_e_*(*t*) and the fix distance, *d*_0_ (in mm), between the magnet and the HES when the shell is closed:

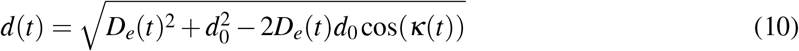

From the calculation of *d*(*t*), we calculated the gape angle *α*(*t*) (in *radian.s*^−1^) as

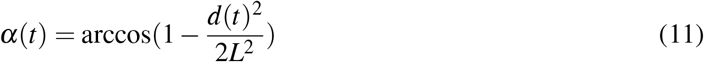

which, for presenting results, were converted in degree by multiplying the right term by 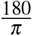

## 3 Results

### 3.1 Mathematical properties of the adductor muscle dynamics model

The opening process, which was formulated in equations [1], [2], and [3] and expressed in [7], takes into account mechanical limits of shell valve opening. The relative opening force is null when the shell is fully open (*α*(*t*) = *α_max_*) and is maximum, equal to 1, when the shell is closed (*α*(*t*) = 0). The equation [7] is at equilibrium if 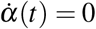, hence if *ε*(*t*)^2^ = *Qp_A_*. It is expressed as:

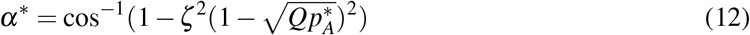

where 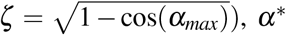, is the gape angle equilibrium value (i.e., resting-state) and 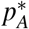 is the equilibrium-state value of the proportion of fibers activated in the smooth part of the adductor muscle. *α*\* is bounded by *α*\* ∈ [cos^−1^(−1), *α_max_*].

The calculation of 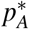 is given by the analytical integration of [5]. The general solution is:

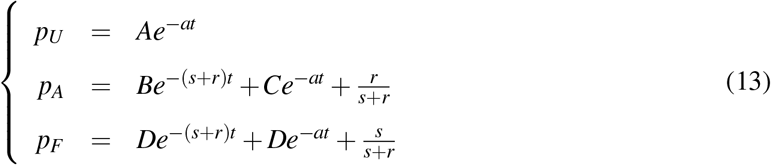

where A, B, C, and D are constants, depending on the parameters and initial values for *p_A_* and *p_F_*. Consequently, the non-trivial stable equilibrium solution does not depend on initial conditions:

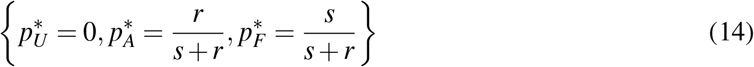

This solution states that, at equilibrium, all muscle fibers are either activated (hence pull valves to close the shell) or deactivated. This consituted a fundamental issue since discrete closing events could not occur any longer in this configuration. In order to solve this problem, the model was revised as:

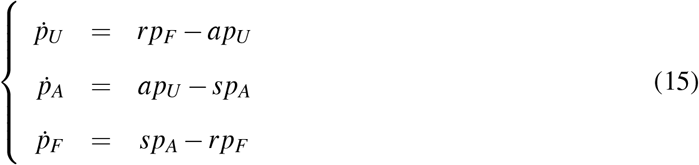

The non-trivial equilibrium of [15] is:

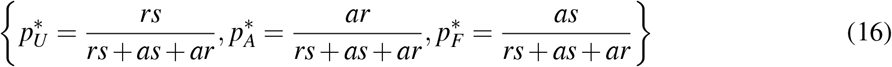

When equation [16] is introduced into equation [7], the opening angle at equilibrium can be determined as:

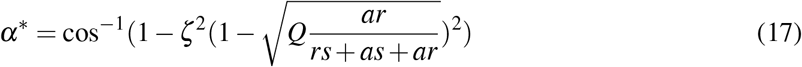

where *α_max_* is measured and 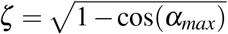. The parameters 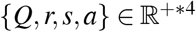 must be identified.

A series of three simulations were performed to illustrate the dynamics simulated by the model (Equations 7, 8 and 15; Figures 2 and 3). Parameters and initial conditions were set as identical for the three simulations ({*a* = 0.0015, *s* = 0.0015, *r* = 0.0007, *K* = 50, *Q* = 0.74}, with r and Q being calculated in such a way that *α_max_* and *α*\* were equal to 30 and 15 degrees respectively). The three numerical simulations were performed at 10 Hz (dt = 0.1 s). The first simulation (Figure 2, upper graph) shows that with constant time duration between two events (fixed at 30 min), the dynamics are characterized by a series of identical closing peaks, all of which have the same intensity. When the period fluctuates randomly (second simulation, Figure 1, lower graph), while parameters remain constant and identical to the first simulation, closure peaks with different intensities characterized the dynamics. Therefore, these changes in peak intensities are only the consequences of the random duration between peaks. Finally, a series of very short, random durations between two closing events were examined (Figure 2). For this third simulation, we set T to fluctuate randomly, but we simulated a series of short closing events occuring between ca. 1.0 and 2.0 h; the expectation of the duration between two events decreased from 30 minutes to 1 minute during this time window. This created dynamics, which tended to deviate from the equilibrium value. All of our simulations showed a slightly larger opening event prior to a return to equilibrium. This is because the model [16] behaves like an over-damped oscillator. The amplitude of this damped oscillation depends on the intensity and duration of the contraction as well as on the speed of return to an equilibrium; it simulates a short-term muscle fatigue which leads to a slightly larger opening angle prior to the return to an equilibrium value during a recovery period.

**Figure 2:**
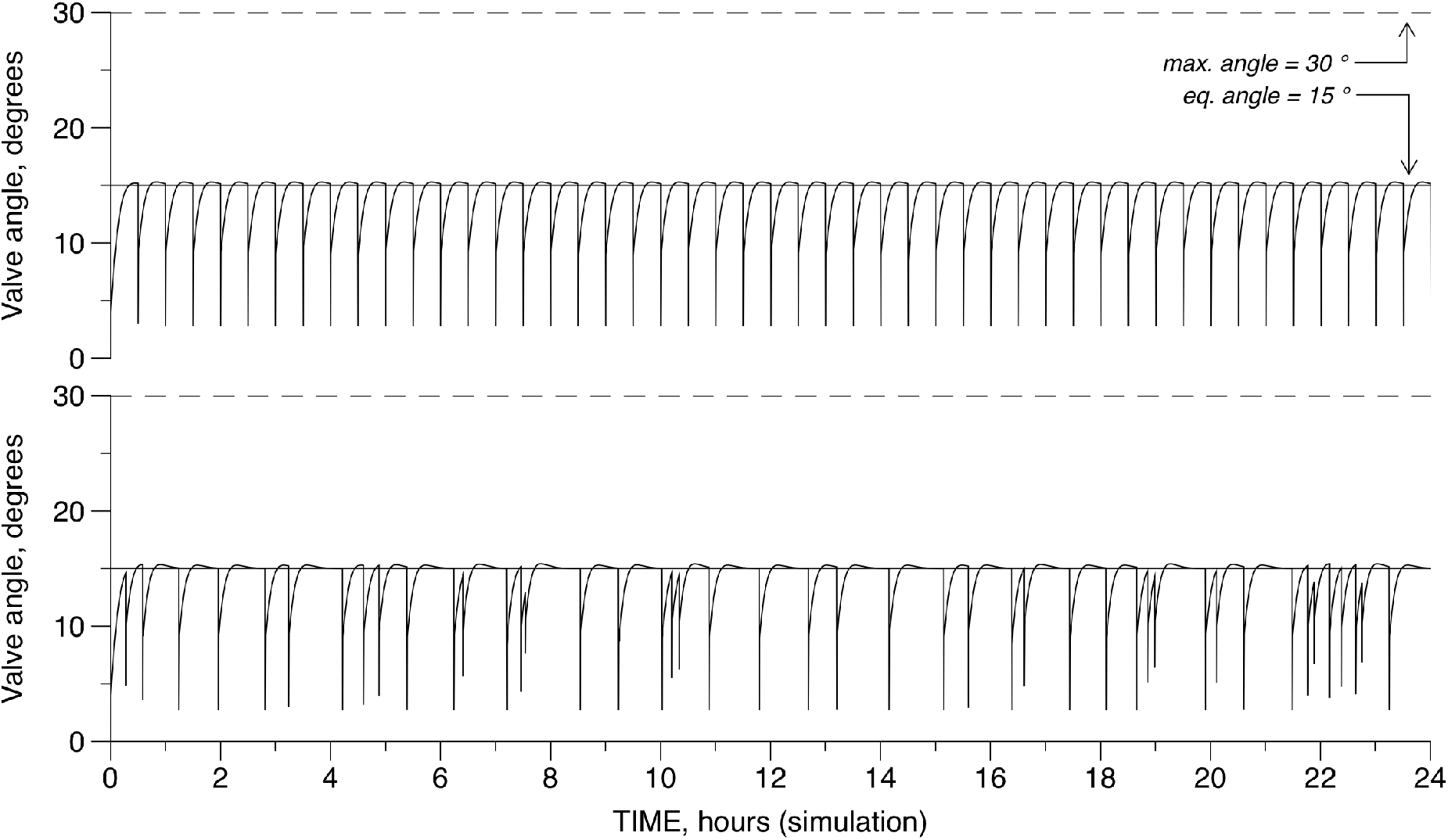
Two simulations performed with the same set of parameters and initial conditions. Parameters were fixed arbitrarily for the three simulations: {*a* = 0.0015, *s* = 0.0015, *r* = 0.0007, *K* = 50, *Q* = 0.74}, with r and Q being calculated in such a way that *α_max_* and *α* * were equal 30 and 15 degrees respectively. Horizontal lines represent the equilibrium, “resting” angle (in degrees), and the maximum opening angle (in degrees). The upper graph shows what data would look like from a model simulating an identical duration between two closing events (30 min). The lower graph presents a data series obtained with the model simulating randomly variable durations (expectation is set equal to 30 min) between two closing events.

**Figure 3:**
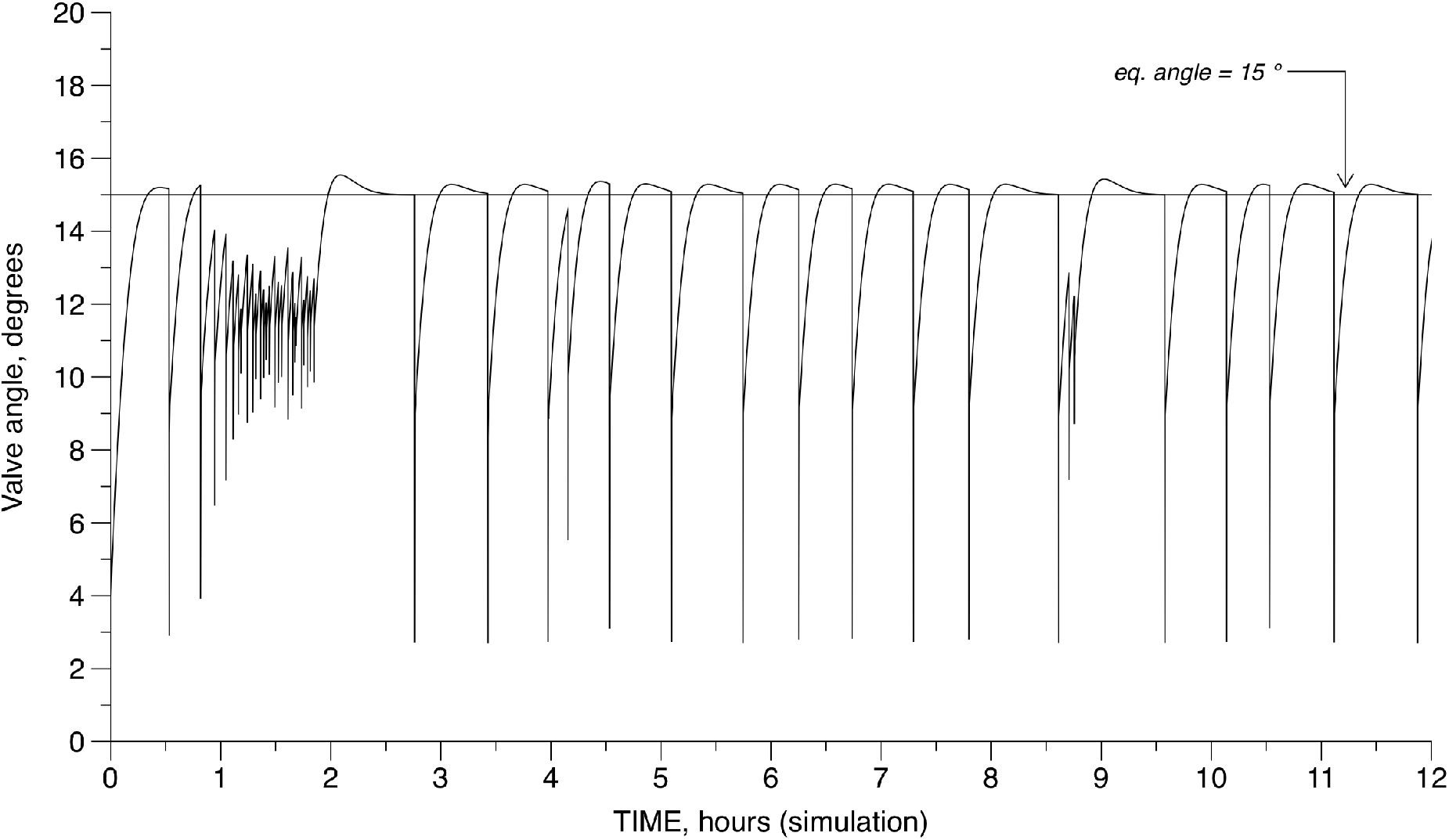
Simulation performed with the same set of parameters and initial conditions as Figure 2. The horizontal line at 15° represents the equilibrium, or “resting” angle. The model simulated randomly variable durations between two closing events, but the expectation decreased from 30 minutes to 1 minutes between ca. 1 and 2 hours, representing a stressful event. It resulted in a period of a sustained effort. The dynamics overpass the equilibrium angle value after events of contraction, proportional to the duration of contraction and the time to return to the equilibrium.

### 3.2 Assessing individual variability

Figure 4 shows two examples of recorded time series both exhibiting discrete events during their continuous dynamics; the upper part corresponds to specimens 4 and the lower part, to specimen 9; specimens 4 and 9 were selected randomly from the experimental group in (Guarini et al. 2020). We kept on using them as examples for more detailed description and discussion in the following sections. The results of the data analysis for all specimen are presented in Table 1; note that data recordings from three specimens, 7, 8 and 10 were not analysed because of excessive variability (in 7 and 8) and errors (in 10) in writing the data file (Guarini et al. 2020).

**Figure 4:**
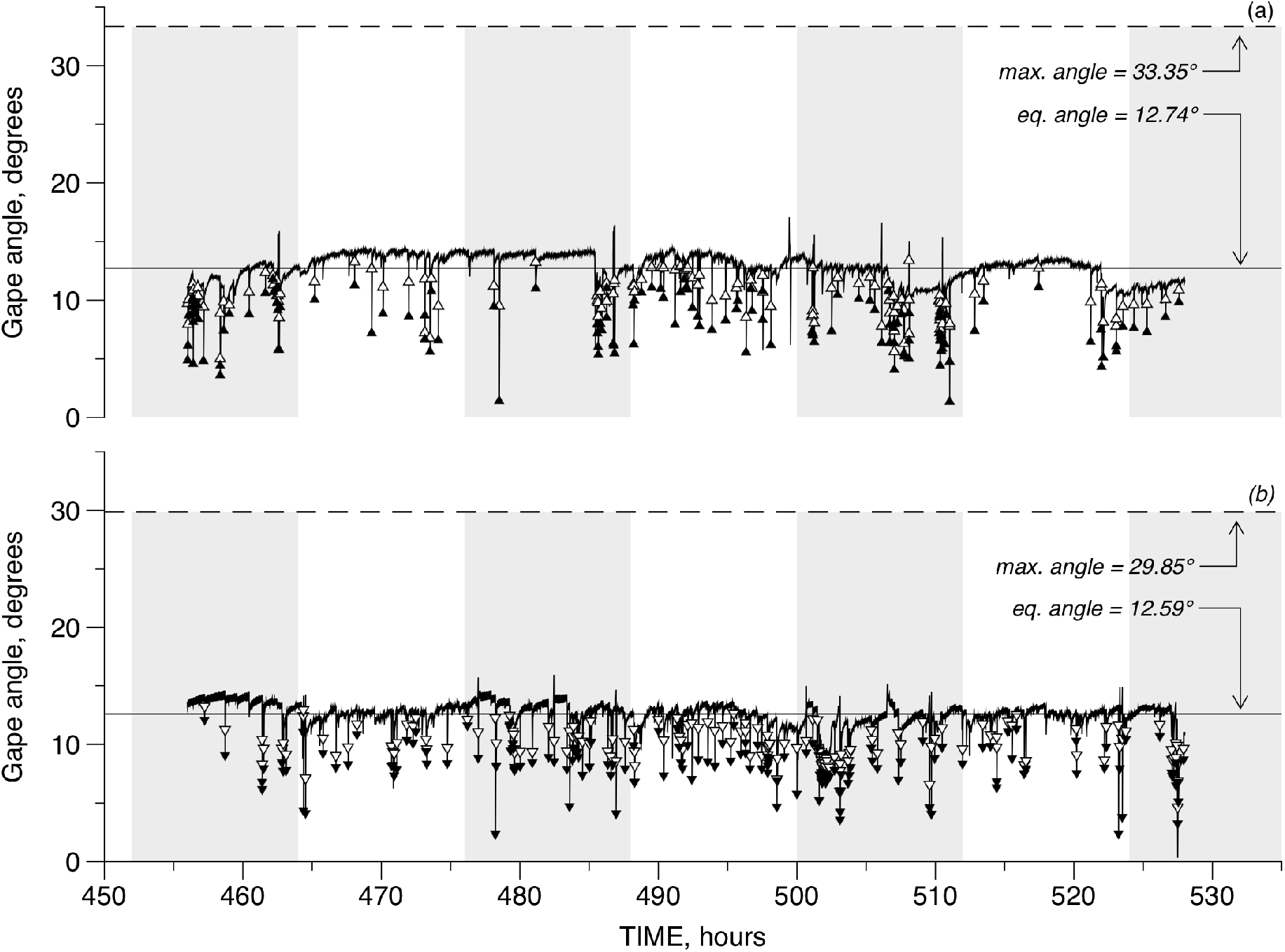
Two different observed HES valve gape time series converted to changes in gape angle for specimens 4 (upper panel, a) and 9 (lower panel, b). The dashed line is the maximum gape angle amplitude, *α_max_*, obtained from calibrations after the adductor muscle was severed. The solid line shows the average gape angle calculated from the estimated time series. The triangles (up for specimen 4, down for specimen 9) indicate where the fitting procedure identified a peak start (empty symbols) and an event peak (filled symbols) for discrete events in each series. The alternating gray blocks indicate the 12 hour dark-light periods in the experimental tanks.

**Table 1.** Parameters of the adductor muscle activity dynamics and the resulting valve opening angle. *α_max_* and *α*\* are in degree, K in degree per second (°.*s*^−1^), Q is a dimensionless scaling factor between opposite forces exercised by the ligament and the adductor muscle. *ρ* is a rate of occurrence between two discrete closing events (*h*^−1^), Peaks represents the number of peaks identified in the time series measured by the Hall-effect valvometer. *χ*^2^ is the value calculated to test differences in the distribution of the duration, *τ*, between two discrete closing events, as observed with the Hall-effect valvometer and as simulated with a pure stochastic process. d.o.f. is the degree of freedom corresponding to the calculation of *χ*^2^, used to compare these values with a critical value obtained from a table, when fixing the p-value.

| Specimen | $\alpha_{max}$ | $\alpha^*$ | K | Q | $\rho$ | Peaks | $\chi^2$ | d.o.f. |
| --- | --- | --- | --- | --- | --- | --- | --- | --- |
| 1 | 29.85 | 14.43 | 14.75 | 0.79 | 0.35 | 417 | 8.53 | 9 |
| 2 | 37.16 | 14.56 | 16.47 | 0.92 | 0.45 | 322 | 16.33 | 9 |
| 3 | 36.28 | 17.85 | 25.07 | 0.76 | 0.36 | 396 | 9.27 | 9 |
| 4 | 33.35 | 13.74 | 15.25 | 0.92 | 0.47 | 307 | 11.09 | 9 |
| 5 | 29.85 | 15.02 | 11.99 | 0.73 | 0.31 | 381 | 15.52 | 9 |
| 6 | 37.16 | 19.85 | 39.93 | 0.63 | 0.08 | 1439 | 16.13 | 9 |
| 7 | *excess noise* |  |  |  |  |  |  |  |
| 8 | *excess noise* |  |  |  |  |  |  |  |
| 9 | 29.85 | 12.63 | 12.67 | 1.01 | 0.38 | 318 | 10.97 | 9 |
| 10 | *data write errors* |  |  |  |  |  |  |  |
| 11 | 36.28 | 13.93 | 17.05 | 0.97 | 0.31 | 382 | 10.41 | 9 |
| 12 | 33.35 | 17.08 | 14.78 | 0.70 | 0.64 | 148 | 9.13 | 9 |

A total of 307 and 318 closing peaks were identified for specimens 4 and 9, respectively. The closing event rate, *ρ* (in *h*^−1^) was estimated to be equal to 0.47 *h*^−1^ (specimen 4) and 0.38 *h*^−1^ (specimen 9). For each of the graphs in figure 4 are presented the observed angle variations estimated from the HES calibration (for each individual), the average angle value (solid horizontal line) and the maximum angle value (dashed horizontal line, estimated from the maximum valve gape distance measured after the adductor muscle was severed). The estimated boudaries of the closing peaks are presented with the starting upper values (unfilled triangles) and the ending lower values (filled triangles) which were estimated using the fitting procedure described in this article. The average angle values were equal to 12.74° and 12.59° for specimens 4 and 9, respectively. The measured maximum opening angle values were 33.35° and 29.85°, respectively (Table 1).

The second step consisted of identifying the parameters {*K, Q*}, from system [16] with transition [17]. We were not able to identify all four parameters {*K, Q, r, s, a*} because there is no quantitative information available about the functioning of the muscle fibers. {*K, Q*} were estimated from equation [17] by fixing {*r,s,a*} at a constant value equal to 0.0015; this was estimated for the 9 available observed time series by minimizing the squared differences between the estimated and corresponding calculated beginning and end values of the peaks, between days 2 and day 6 (Table 1). A direct search algorithm (Nelder and Mead, 1965) was used to do the minimization of the least squares criterion (Table 1). From these parameter estimates, *α*\* was calculated (equation 17).α*, which corresponds to the equilibrium value toward which the continuous model (equation 7) converges asymptotically, is different from the average angle estimates, which is calculated from all estimated angle of the observed data series. Figure 4 shows the simulations regarding the observations, after being optimized to the identified peaks. Parameters, K and Q, were equal to 15.25°.*s*^−1^ and 0.92 °.*s*^−1^ for specimen 4, respectively (Figure 4, upper graph), and were 12.67°.*s*^−1^ and 1.01 °.*s*^−1^ for specimen 9 (Figure 4, lower graph), respectively. For specimen 4, the equilibrium angle (*α*\* = 13.74°) of the simulated series surpassed frequently the average opening angle value of the observed series. However, for specimen 9, the equilibrium angle (*α*\* = 12.63°) of the simulated series remained very close to the average opening value of the observed series. For both example specimens, periods of high frequency occurrences of closing events (at 455, 485 and 505 hours, for specimen 4 and at 500 hours for specimen 9) were associated with lower valve gape openings. In the case of specimen 4, a period of lower valve gape opening, at 520 hours, without a higher frequency was observed.

Figure 5 shows that the simulated results approximated the observation data, but closure peak estimates were often smaller than the overall observations. In addition, the simulations showed more regularity in closure peak amplitude than the observations for both specimens. The simulations represented well the decreases of valve gape angle amplitude and maximum values when the peak occurrence frequency increased.

**Figure 5:**
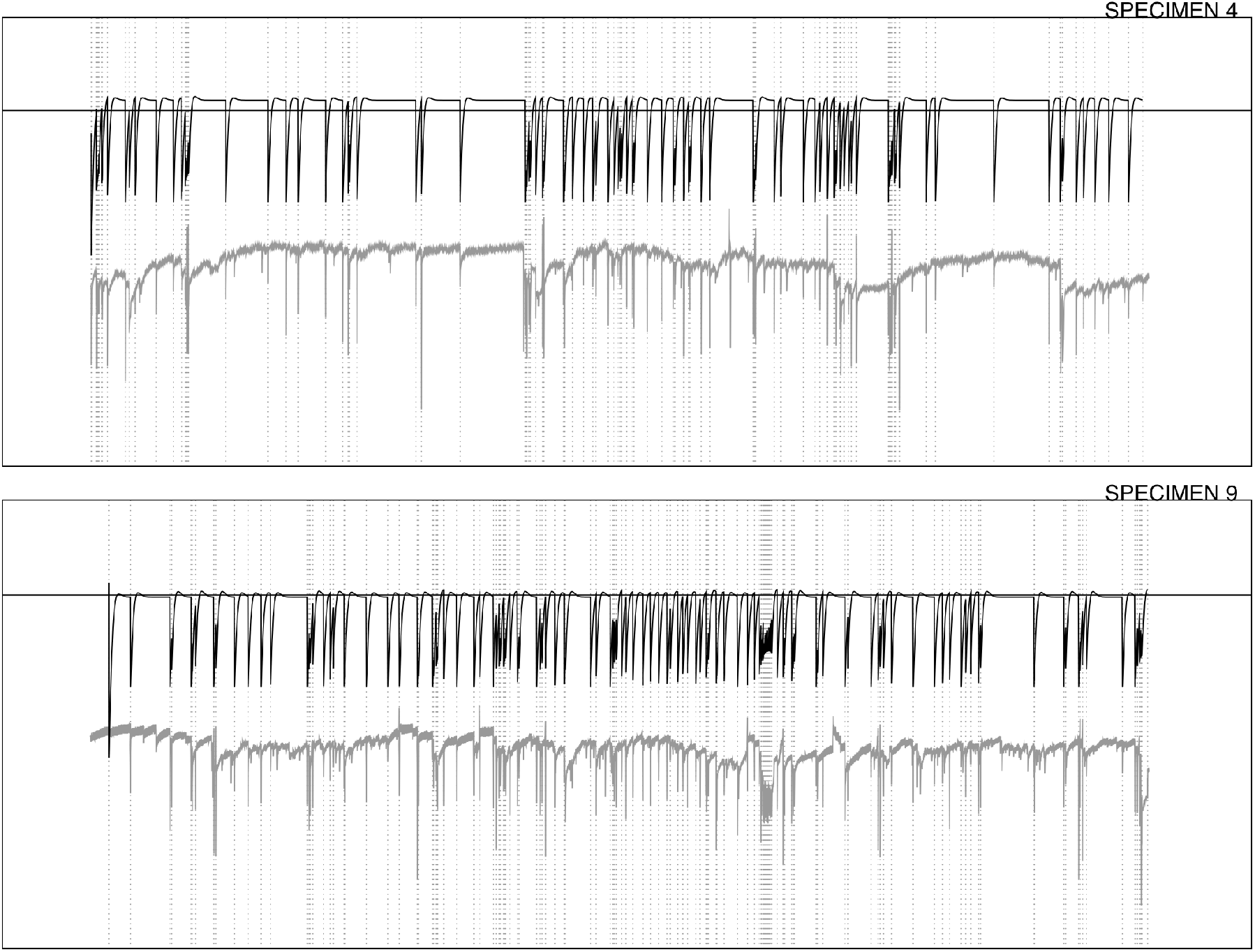
Qualitative comparison of the simulated and observed valve gape dynamics for specimens 4 (upper panel) and 9 (lower panel). The black lines are the simulated series for the respective individuals. Below this are the data series in light gray. Simulated series were performed by optimization of an ordinary least square criterion, identifying parameters Q and K (a, r, s being fixed at 0.0015). The optimization was performed on peaks occurrence only (their placement is indicated by the vertical dashed lines). The horizontal, solid lines indicate the resting gape angle calculated from parameter values(*α_max_*), characteristic of the equilibrium state in the adductor muscular fibers.

Parameter values in Table 1 indicate the estimated angles of *α_max_* can vary from 29.85° (specimen 1) to 37.16° (specimen 6), with an average value of 33.68° among the nine individuals.

The equilibrium opening angle *α*\* varied from 12.63° (specimen 9) to 19.85 ° (specimen 6), with an average value of 15.45°. The number of identified closing peaks and related *ρ* values varied from 148 peaks and 0.64 *h*^−1^ (specimen 12) to 1439 peaks and 0.08 *h*^−1^ (specimen 6). The values for specimens 12 and 6 differed greatly from the other seven specimens which had average values of 360 peaks and *ρ* = 0.38 *h*^−1^, respectively. In addition, for these seven specimens, the values of K varied from 11.99 deg.*s*^−1^ (specimen 5) to 25.07 deg.*s*^−1^ (specimen 3), with an average value of 16.18 deg.*s*^−1^. For the scaling factor Q, the values varied from 0.63 (specimen 6) to 1.01 (specimen 9), with an average value of 0.82.

In our approach, the occurrence of discrete events was assumed to be determined externally; this means that they were not triggered by the state of the continuous dynamics. The discrete dynamics can then be considered as a series of transitions between continuous periods. The occurrences of these transitions were tested to determine if they were spaced randomly. The observed time series were thus compared to a stochastic process defined as null models. These null models were designed to simulate random series of discrete events with an average occurrence rate, 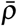 (in *h*^−1^), calculated from the corresponding observations. The time *τ* (in h) between two discrete events was calculated as:

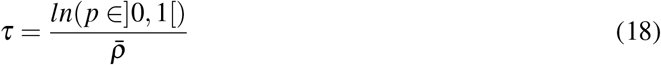

where *p* ∈] 0, 1 [is a uniformly distributed probability; consequently, *τ* follows the exponential law, *f* (*τ, ρ*). Comparisons between the observed and null distributions were made with a Pearson’s *χ*^2^ test. The degree of freedom was fixed to 9 (optimal number of classes - 1) and the first-type error was fixed at 0.05. *χ*^2^ values were found to vary between 8.53 (specimen 1) and 16.33 (specimen 2). With a critical value 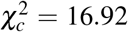, the distributions of the time intervals between any two events could not be distinguished from pure stochastic processes with a confidence of 0.95. Specimens 2, 5, and 6, for which *χ*^2^ values approached 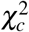, all had short periods of rapid shell clap sequences that induce an increased number of short duration closures in the series.

## 4 Discussion

### 4.1 The opening angle: a non-measurable, yet indispensable, variable

The angle *α* is the central variable used in our study. It is calculated from the valve gape distance which is estimated from the HES protocol. It is used in valvometry studies (e.g. Comeau et al. 2012) because it standardizes the dimensions associated with the valve opening process. Thus the valve gape angle, on the contrary of the valve gape distance, is assumed to be independent from the shell size, which may increase over long experimental time.

The variations of the opening angle, *α*, were expressed as a function of the forces applied to the shell valves. In our study, we have consistently taken into account, in all our estimate calculations, the fact that bivalve shells rotate around their hinge axis. These angular forces include the passive action of the ligament and the active effort produced by the adductor muscle. The force produced by the contraction of the muscle is a function of the physiological energy spent by the organism, hence depends on its physiological condition. In our modelling framework, parameter identification depends on the estimated values of the opening angles *α* with a particular realization, *α_max_*. However, the opening angle, *α*, is not measurable; it is estimated from valvometry data, acquired by the HES. The opening angle is a geometric construction of a triangle and thus defined by the angle between three vertices of the shell. One vertex is located near the umbo, on the virtual axis of rotation for the valves. The two other vertices are both determined by the positions of the HES. One of these two positions is mobile (position of the HES when it moves) and the other is a fix point, corresponding to the position of the HES when the shell is closed (Figure 1A). The three vertices define an isoceles triangle, because two vertices are defined by the HES positions. In all our calculations, the quality of the resulting *α* estimates depends on the accuracy of the position of the HES regarding to the umbo and the magnet; unbiased and accurate estimates are achieved when the HES and the magnet are perfectly aligned when the valves are closed. Therefore, errors in the positioning of the HES and the magnet and the measurements of these positions generate biases in calculations of *α*, the forces applied, and related estimators. As there were no means to verify this assumption *a posteriori*, all values presented here depend on the respect of this alignment during the preparation of the experiment. We recommend for future study to equip each individual with two different HES sensors, in order to investigate possible problems of positioning and functioning.

### 4.2 Modeling the opening angle dynamics

There are many studies of the adductor muscle in bivalves, but we did not find any quantitative studies that linked muscle activity and angle opening dynamics. In our study, we chose to simulate the dynamics of the opening angle as a function of the variations in smooth muscle activity. Two parameters of the opening angle dynamics in equation [16], K and Q, were estimated by optimization (minimizing the distance between simulations and experimental observations). K is an angular velocity (.*s*^−1^), while Q is a (unitless) scaling factor between the opposite forces produced by the ligament and the adductor muscle.

Based on our analysis of the muscle contraction model, we found that the continuous dynamics are characterized by a constant convergence to a non-trivial stable equilibrium state, which depends on the values of the parameters Q,a,r,s and on the maximum opening angle *α_max_*. This equilibrium is then understood as the balance between the passive opening force (*i.e*., produced by the elongation of the ligament, and an active force (*i.e*., produced by the contraction of the adductor muscle. Under the equilibrium state condition, there is also a balance between the processes of activation, de-activation, and recuperation of the muscle fibers. The equilibrium opening angle therefore depends on the capacities of an individual scallop to maintain a fraction of the muscle fibers activated. If the individual cannot maintain this condition, the shell opens to reach the maximum gape value, close to *α_max_*.

The model presented was designed to simulate the gape variations of the valves and explain the dynamics of shell opening in terms of its mechanical properties. However, the biophysical parameters of the adductor muscle submodel remain unidentifiable; by this we meant that their values cannot be estimated by optimization only without acquiring specific information regarding the activity and physiological state of the organism. Therefore, we fixed the parameters {*a, r, s*} in order to estimate the two other parameters (K and Q) of the main differential equation. The parameters {*a, r, s*}, however, are supposed to change with environmental conditions (e.g. temperature, oxygen concentration …) and physiological conditions (MacDonald et al. 2006). In our model framework then, each parameter can be replaced by a function describing environmental variables that may affect the processes that they control. If ancillary measurements and complementary experiments were to become available, the model framework here would be well-suited to identifying and quantifying the related processes.

Our model also accounted for discrete events. One of the major issues addressed in our development was the revision of the muscle dynamics submodel to allow discrete events to occur (see section 3.1). This is an important goal for any applications of our framework because aquatic organisms are subject to many different types of short, transient natural and anthropogenic events (*e.g*. ambient noise, turbidity plumes, salinity drops, toxic algal blooms …) that interact with longer, seasonal environmental and biological cycles. The formulation originally developed by Liu et al. (2002) as a generic model for human muscles was unable to simulate the occurrence of discrete events in mollusk valves dynamics once equilibrium states were achieved.

Discrete closing events were considered to be the result of an external control. They could be determined internally as a function of the state variables of the continuous model. We found that the time between two discrete events cannot be differentiated from a random process, in all our observations. This was not the case for the amplitude of the peaks, which did not follow this pattern. These remarks suggest that we reached some limits in our study to understand the valves dynamics physiologically. On the one hand, some estimators (like amplitude) could have been determined more accurately, but on the other hand, the functioning of the smooth adductor muscle is complex, and may involve interactions between several processes, producing a random-like pattern. In particular, the smooth adductor muscle is able to perform two kinds of contractions, tonic and phasic. As a result, sustained effort with low energy consumption is achieved, maintaining a capacity to perform rapid, dynamic contractions with controlled intensity (Chandler 2006). By linking the muscle activity to nervous stimuli and by establishing energetic cost within a context of adaptive strategy (Guderley and Portner 2010), the capacity of an organism to react internal conditions, as well as to its surrounding environment, could be determined. Interestingly, the energetic cost of valve movement (which can be estimated from the forces) has never been treated in any bioenergy budget model. Energy management, from uptakes to utilization, should be investigated by integrating different functions and behaviors of the scallop (MacDonald et al. 2006).

The model still only partially reproduces the variability of the observed hybrid dynamics. A more comprehensive eco-physiological model which includes adequate observations about muscle activity and other physiological functions, as well as corresponding stimulations of the nervous system, is now required to advance further. In particular, the perceived stochastic nature of discrete closing events, their control (internal and external) and their energetic cost function of the physiological state of the organism should all be determined.

### 4.3 Smooth versus striated muscle dynamics and micro-contractions

Bayliss (1930) and Wilkens (1981) first described differential dynamics of the smooth and striated parts of the adductor muscle. The nervous system that controls the adductor muscle has the task of managing both the smooth muscle fibers for sustained effort and striated muscle fibers for fast transient responses (Wilkens 2006). In addition to single closing events, fast shell clappings over short periods, to swim, rotate or bury in the sediment, are related to the activity of the striated muscle fibers (Wilkens, 2006).

However, a formalization of these reflex-contractions and its integration within the continous dynamic regime appeared to be difficult to achieve. The first problem was that it cannot be simulated by the same system of equations as [15] because reflex-type dynamics cannot not be represented by such processes. A model that includes both contraction and relaxation cycles is required. There are many existing functions which can fulfill these properties, and they are generally represented by second order differential equations. For example:

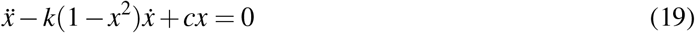

which can be decomposed, introducing *y* = *x* and 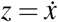 as a system of two first order equations:

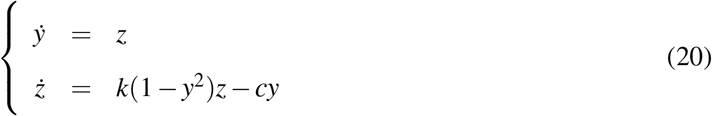

where k and c are two parameters, and y the state of muscle (contraction or relaxation) and z represent the changing rate. The system describing variations of y and z simulate an oscillator, and the resulting force of the striated muscle is described by an exponential function of y (Huxley and Simmons 1971).

A second problem is that the transition between the continuous dynamics and the reflex-type dynamics has to be formalized. Particularly, a system which simulates active relaxation of the smooth part of the adductor muscle should be formulated (Wilkens 1981). For example:

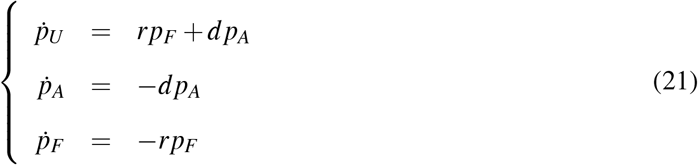

where d represents an active relaxation rate in *t*–1. Such a transition would have to be explicitly controlled among all other possible controls of the valve gape dynamic processes, by the nervous system. Nervous system modelling is a prolific field with many hypotheses about stimulation and signal propagation (Izhikevich, 2010) which lead to a muscle contraction. Therefore, the implementation of a nervous control in our model, even if possible theoretically, would require experiments with induced stimulations, to provide relevant insights into the valve gape dynamics.

The third problem is to determine the duration of the reflex-type dynamics, hence to control the dynamics by the individual energy consumption, since in its present from, the model [20] can sustain oscillations without limits.

Our last remarks concerns the observed high frequency, small-scale variability in muscle contractions, superimposed onto the global dynamics. In Figure 6, the upper graphs for the two specimen 4 and 9, taken as examples, show global gape angular velocity, while the lower graphs show the detailed small-scale variability around the null velocity (velocity = 0.0 ± 0.7 °.*s*^−1^). The global pattern reveals peaks with average velocities close to the values determined for parameter K; the average velocity value for specimen 4 was slightly higher than that for specimen 9 (ca. 15 °.*s*^−1^ vs. ca. 12 °.*s*^−1^, respectively). In contrast, the small-scale variability (lower figures for each specimen) differed markedly from the global pattern and from one specimen to the other; particular patterns of intervalve micro-contractions were observed, which are consistent with micro-closure events aloready observed in other studied (Comeau et al. 2019). These events were hypothesized to be caused by irritating substances present in the pallial cavity (Galtsoff 1964), such as the presence of toxic algae. In general, micro-contractions are usually attributed to external factors (Comeau et al. 2019). In our experimental system, water input quality was controlled during the measurement period. Thus, without excluding causes that were unintented and not monitored, we hypothesize that these micro-contractions were produced in response to internal factors and they may reveal physiological processes, *i.e*., calcification events along the shell edge. It will be important for these micro-contractions to be characterized comprehensively in future studies.

**Figure 6:**
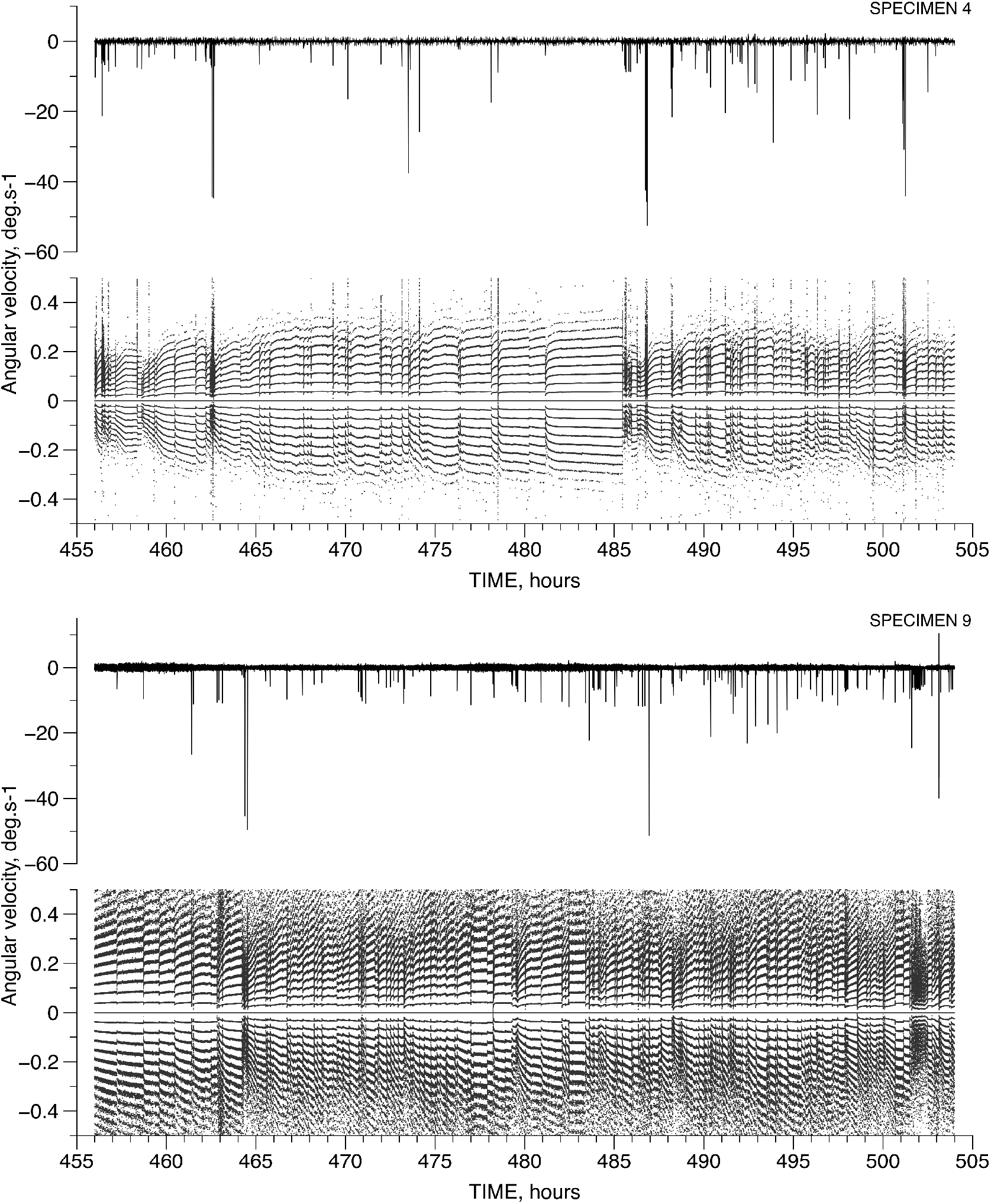
High frequency, small amplitude dynamics of inter-valve closures and openings compared to a global pattern. These variations were estimated according to angular velocity variations, which showed changes in acceleration. These changes were interpreted as periods of rapid contraction and slower relaxation. The upper two graphs represent results for specimen 4 and the lower two graphs, for specimen 9. For each series of two graphs, the upper one represents the global pattern (on the full range of velocity) and the lower one, the small-scale variability around the null velocity (0.0 ± 0.7 °.*s*^−1^). More discrete contractions and a particular pattern can be observed at small scale compared to the global scale; they are therefore characterized as micro-contractions superimposed on the macro-scale pattern and may reflect physiological processes.

## 5 Conclusions

The model developed here for *Pecten maximus* is an important step forward for the prediction of valve opening dynamics from forces generated by the smooth adductor muscle. The model simulates valve dynamics described by a series of continuous convergences to an equilibrium state, interspersed with random discrete closing events. This outcome provides a perspective to exploit valve movements as indicators of the health of bivalve organisms. It remains to consider how to quantify the neurophysiological control of muscle activity in such a way that external and internal stimuli could modulate the activation or deactivation of the adductor muscle fibers. Model expansion should also integrate chemical cues for the nervous system controlling the adductor muscle. This will permit combining the biophysical system with estimates of the physiological capacities of the organisms. The biophysical model should then be enriched with biochemical processes using the initial model formulated by Wexler et al. (1997) with Ding et al. (2000) modifications to account for the physiological state of the adductor muscle. However, further model development will require ancillary *ad hoc* measurements in order to identify parameters and processes that control the dynamics of resulting systems. Therefore, two research paths can be identified with two different goals and perspectives. First, laboratory experimentation associated with new model development should increase our understanding of ecophysiological mechanisms. Second, *in situ* monitoring of bivalve behavior can be linked more closely to the model describing valve gape dynamics through data assimilation methods. This will permit optimal forecasts in real time, improving environmental impact and monitoring programs in return.

## Data and Code accessibility

This text is omitted for the preprint.

## Acknowledgements

This text is omitted for the preprint.

## Competing interests

This text is omitted for the preprint.

## Notes

### Competing Interest Statement

The authors have declared no competing interest.

## References

Bayliss LE, Boyland E, Ritchie AD. 1930 The adductor mechanism of Pecten. Proc. R. Soc. B, 106, 363–376.

Böl M, Stark H, Schilling N. 2011 On a phenomenological model for fatigue effects in skeletal muscles. J. Theor. Biol., 281, 122–132.

Chantler PD. 2006 Scallop Adductor Muscles: Structure and Function. In: SE Shumway (ed). Scallops: Biology, Ecology and Aquaculture. Developments in Aquaculture and Fisheries Science, 35, 229–316.

Clements JC, Comeau LA. 2019 Use of High-Frequency Non-invasive Electromagnetic Biosensors to Detect Ocean Acidification Effects on Shellfish Behavior. J. Shellfish Res., 38(3), 811–818.

Comeau LA, Mayrand E, Mallet A. 2012 Winter quiescence and spring awakening of the Eastern oyster *Crassostrea virginica* at its northernmost distribution limit. Mar. Biol., 159, 2269–2279.

Comeau LA, Babarro JMF, Longa A, Padin XA. 2018 Valve-gaping behavior of raft-cultivated mussels in the Rìa de Arousa, Spain. Aquaculture Reports, 9, 68–73.

Comeau LA, Babarro JMF, Riobo P, Scarratt M, Starr M, Tremblay R. 2019 PSP-producing dinoflagellate *Alexandrium minutum* induces valve microclosures in the mussel *Mytilus galloprovincialis*. Aquaculture, 500, 407–413.

Ding J, Wexler A, Binder-Macleod S. 2000 A predictive model of fatigue in human skeletal muscles. J. App. Phys., 89, 1322–1332.

Gainey LF, Shumway SE. 1988 A compendium of the responses of bivalve mollusks to toxic dinoflagellates. J. Shellfish Res., 7, 623–628.

Galtsoff PS. 1964 The American oyster *Crassostrea virginica* Gmelin. U.S. Fish Wildlife Serv. Fish. Bull., 64, 1–480.

Guarini JM, Coston-Guarini J, Comeau, LA. 2020 Calibrating Hall-Effect valvometers accounting for electromagnetic properties of the sensor and dynamic geometry of the bivalves shell. BioRχiv. doi: https://doi.org/10.1101/2020.12.20.423648

Guderley H, Portner HO. 2010 Metabolic power budgeting and adaptive strategies in zoology: examples from scallops and fish. Can. J. Zool., 88, 753–763.

Huxley AF, Simmons RM. 1971 Proposed mechanism of force generation in striated muscle. Nature, 233, 533–538.

Izhikevich EM. 2010 Dynamical Systems in Neuroscience:The Geometry of Excitability and Bursting. Computational Neuroscience Series. The MIT Press Cambridge, Massachusetts; London, England. 464 pages.

Kramer KJM, Jenner HA, Dick de Zwart D. 1989 The valve movement response of mussels: a tool in biological monitoring. Hydrobiologia, 188/189, 433–443.

Liu JZ, Brown RW, Guang HY. 2002 A Dynamical Model of Muscle Activation, Fatigue, and Recovery. Biophys. J., 82, 2344–2359.

Looft MJ, Herkert N, Frey-Law L. 2018 Modification of a three-compartment muscle fatigue model to predict peak torque decline during intermittent tasks. J. Biomechanics, 77, 16–25.

MacDonald BA, Bricelj VM, Shumway SE. 2006 Physiology: Energy Acquisition and Utilisation. In: SE Shumway (ed). Scallops: Biology, Ecology and Aquaculture. Developments in Aquaculture and Fisheries Science, 35, 417–492.

Marceau F. 1909 Recherche sur la morphologie, et l’histologie, et la physiologie compares des muscles adducteurs des mollusques acéphales. Arch. Zool. Exp. Gen., 5(2), 295–469.

Nagai K, Honjo T, Go J, Yamashita H, Oh SJ. 2006 Detecting the shellfish killer *Heterocapsa circular-isquama* (Dinophyceae) by measuring bivalve valve activity with a Hall element sensor. Aquaculture, 255, 395–401.

Nelder VA, Mead R. 1965 A Simplex method for function minimization. Computer J., 7, 308–13.

Payton L, Sow M, Massabuau J-C, Ciret P, Tran D. 2017 How annual course of photoperiod shapes seasonal behaviorof diploid and triploid oysters, *Crassostrea gigas*. PLoS ONE, 12(10), e0185918.

Rao KP. 1954 Tidal rhythmicity of rate of water propulsion in *Mytilus californianus* and its modifiability by transplantation. Biol. Bull., 43, 283–293.

Redmond KJ, Berry M, Pampanin DM, Andersen OK. 2017 Valve gape behaviour of mussels (*Mytilus edulis*) exposed to dispersed crude oil as an environmental monitoring endpoint. Mar. Poll. Bull., 117, 330–339.

Wexler A, Ding J, Binder-Macleod S. 1997 A mathematical model that predicts skeletal muscle force. IEEE Transactions on Biomedical Engineering, 44, 337–348.

Wilkens LA. 1981 Neurobiology of the scallop. I. Starfish-mediated escape behaviours. Proc. R. Soc. of London B, 211, 341–372.

Wilkens LA. 2006 Neurobiology and behaviour of the scallop. In: SE Shumway (ed). Scallops: Biology, Ecology and Aquaculture. Developments in Aquaculture and Fisheries Science, 35, 317–356.

